# Validation of Davson’s equation in patients suffering from idiopathic Normal Pressure Hydrocephalus

**DOI:** 10.1101/268698

**Authors:** Afroditi-Despina Lalou, Virginia Levrini, Matthew Garnett, Eva Nabbanja, Dong-Joo Kim, Laurent Gergele, Anna Bjornson, Zofia Czosnyka, Marek Czosnyka

## Abstract

**Introduction:** The so called Davson’s equation relates baseline intracranial pressure (ICP) to resistance to cerebrospinal fluid outflow (Rout), formation of cerebrospinal fluid (I_f_) and sagittal sinus pressure (P_SS_) There is a controversy over whether this fundamental equation is applicable in patients with normal pressure hydrocephalus (NPH). We investigated the relationship between Rout and ICP and also other compensatory, clinical and demographic parameters in NPH patients.

**Method:** We carried out a retrospective study of 229 patients with primary NPH who had undergone constant-rate infusion studies in our hospital. Data was recorded and processed using ICM+ software. Relationships between variables were sought by calculating Pearson product correlation coefficients and p values.

**Results:** We found a significant, albeit weak, relationship between ICP and Rout (R=0.17, p=0.0049), Rout and peak-to-peak amplitude of ICP (AMP) (R=0.27, p=3.577e−05) and Rout and age (R=0.16, p=0.01306).

**Conclusions:** The relationship found between ICP and Rout provides indirect evidence to support disturbed Cerebrospinal fluid circulation as a key factor in disturbed CSF dynamics in NPH. Weak correlation may indicate that other factors: variable Pss and formation of CSF outflow contribute heavily to linear model expressed by Davson’s equation.

## Introduction

Normal pressure hydrocephalus (NPH) is a syndrome characterised by ventriculomegaly and the clinical triad of gait ataxia, dementia and urinary incontinence [1, 16, 24]. Despite this syndrome being first described over 50 years ago and presenting an opportunity for symptom improvement following shunt surgery (it is also described as an only kind of reversible dementia), we still know little of its pathophysiology and hydrodynamics [5, 6, 24, 28].

The disturbance of cerebral blood flow and autoregulation periventricularly, with increased intensity proximal to the ventricles has been shown[22]. A variety of processes, possibly including tissue distortion, accumulation of vasoactive and toxic substances, watershed ischaemia and damage in the vasculature could contribute to the pathophysiology and clinical presentation of NPH[[11, 12, 22, 24]. The cause of ventricular enlargement in idiopathic NPH (iNPH) is not as clear. It is thought to be due to a hydrodynamic deficit, although whether this is due to “cisternal block” or aqueduct stenosis or as a problem of cerebrospinal fluid (CSF) absorption due to arachnoid cell hyperplasia and leptomeningeal fibrosis, as well as other degenerative changes, is still uncertain[4–6, 13, 25].

The so called “Davson’s equation” is a fundamental equation in the description of CSF hydrodynamics in physiological individuals^7^. It relates baseline intracranial pressure (ICP) to resistance to cerebrospinal fluid outflow (Rout), formation of cerebrospinal fluid (I_f_) and sagittal sinus pressure (P_SS_), as shown below[24]:

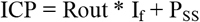

This equation is valid if ICP is greater than Pss. Below Pss, ICP may have any value.

It has been suggested that a key deficit in NPH is a disturbance in the Rout variable and treatment for NPH, shunt surgery, is aimed at manipulating Rout[7–11, 24]. A large Dutch study found that a positive outcome of shunt surgery, as measured by gait improvement on the NPH scale and improvement of dementia measured using the Modified Rankin Scale, can be predicted by a Rout of 18mmHg/ml/min or more[7]. Other studies found significant improvement in clinical outcomes following shunting using lower Rout cut-offs ranging from 13-16mmHg/ml/min[8–10] and a 2015 meta-analysis on Rout thresholds for predicting shunt responsiveness concluded that a Rout of 12mmHg/ml/min is most appropriate for accurate prediction[23]. Nevertheless, the latest multi-centre European study on NPH found that Rout has no correlation to clinical outcome and should not be used to exclude patients from treatmen[29]. Additionally, a 16-patient study, using constant-rate infusion tests, found no relationship between Rout and ICP measured during overnight monitoring[14]. This has ignited controversy over whether Rout is implicated in NPH and whether Davson’s fundamental equation is applicable to patients with NPH.

Davson’s equation (Figure 1) can be positively validated in individual cases. The relationship between variable infusion rate and observed pressure is always linear [15, 24]:

**Figure 1:**
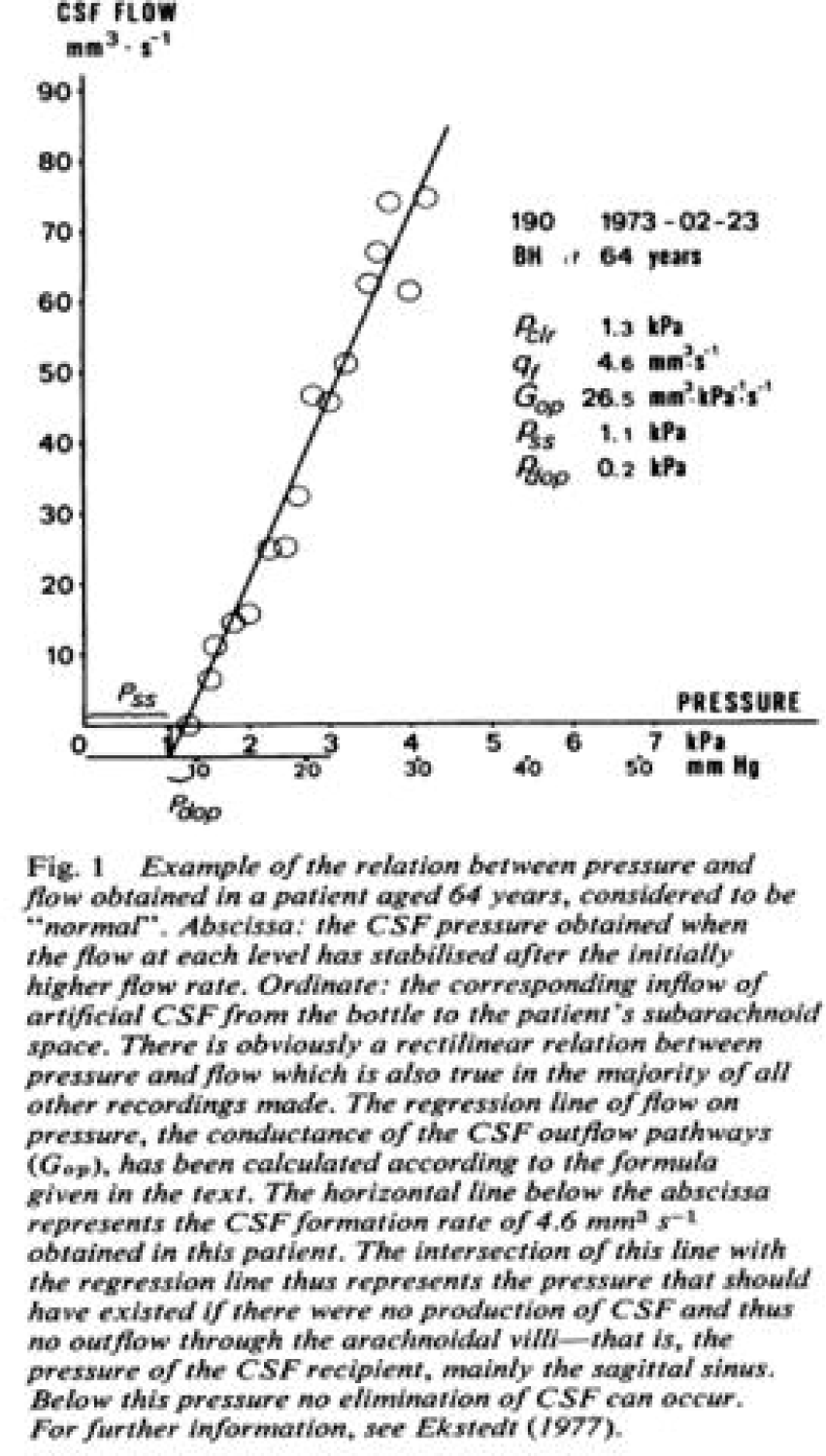
Davson’s equation; The relationship between pressure andfloi,v in a healthy, 64-year-old patient. Reproducedfi,0111 Ekstedt (1977)^15^.

This study investigates the relationship between ICP and Rout between patients in order to ascertain the validity of applying Davson’s equation to NPH patients. In addition, the association between Rout and pulse amplitude of ICP, age, sex, aetiology of patients, etc., has been scrutinized

## Material and Methods

### Patient group

All patients were retrospectively recruited and had a working diagnosis of possible idiopathic NPH (iNPH), with documented radiological evidence of ventriculomegaly on CT and/or on MRI scans, baseline CSF pressure below 18mmHg and at least two of the three cardinal symptoms of NPH (gait disturbance, cognitive impairment, urinary incontinence) including gait disturbance. They attended Cambridge University Hospital Hydrocephalus Clinic between 2009-2013 and an infusion test was indicated as part of their work-up and diagnostic criteria according to hospital guidelines and in line with National Institute for Health and Clinical Excellence (NICE) guidelines[25]. All subjects were provided with an information leaflet and signed individual consent forms. There is some data overlap between this study and previous publications from our databases[13, 19, 23].

### Infusion test

Access was gained via lumbar puncture at the intervertebral space L4-L5 using local anaesthesia or via a previously placed Ommaya reservoir (this is a reservoir placed under the scalp alone, with no associated shunt) with the patient lying on their side. Connection of a fluid filled pressure transducer (Edwards Lifesciences™) and pressure amplifier (Spiegelberg or Philips) to the LP needle allowed for datapoint recording at a frequency of 30-100Hz, with following processing by ICM+ (University of Cambridge Enterprise Ltd)[26]. Once adequate CSF pressure and arterial waveforms readings had been achieved, baseline measurements were taken for 10 minutes, followed by infusion of Hartmann’s solution at 1.5ml/min until the ICP had plateaued for 5-10minutes. As a safety measure, if ICP increased to 40mmHg or above the infusion was stopped. The total duration of the infusion tests was approximately 30 to 45 minutes. Once the infusion test was concluded, a tap test withdrawal of 30 – 50 mls of CSF was carried out prior to removal of LP needle and the patient was kept in hospital for observation for 4 hours. An example of a typical infusion test recording and analysis can be found in Figure 2.

**Figure 2:**
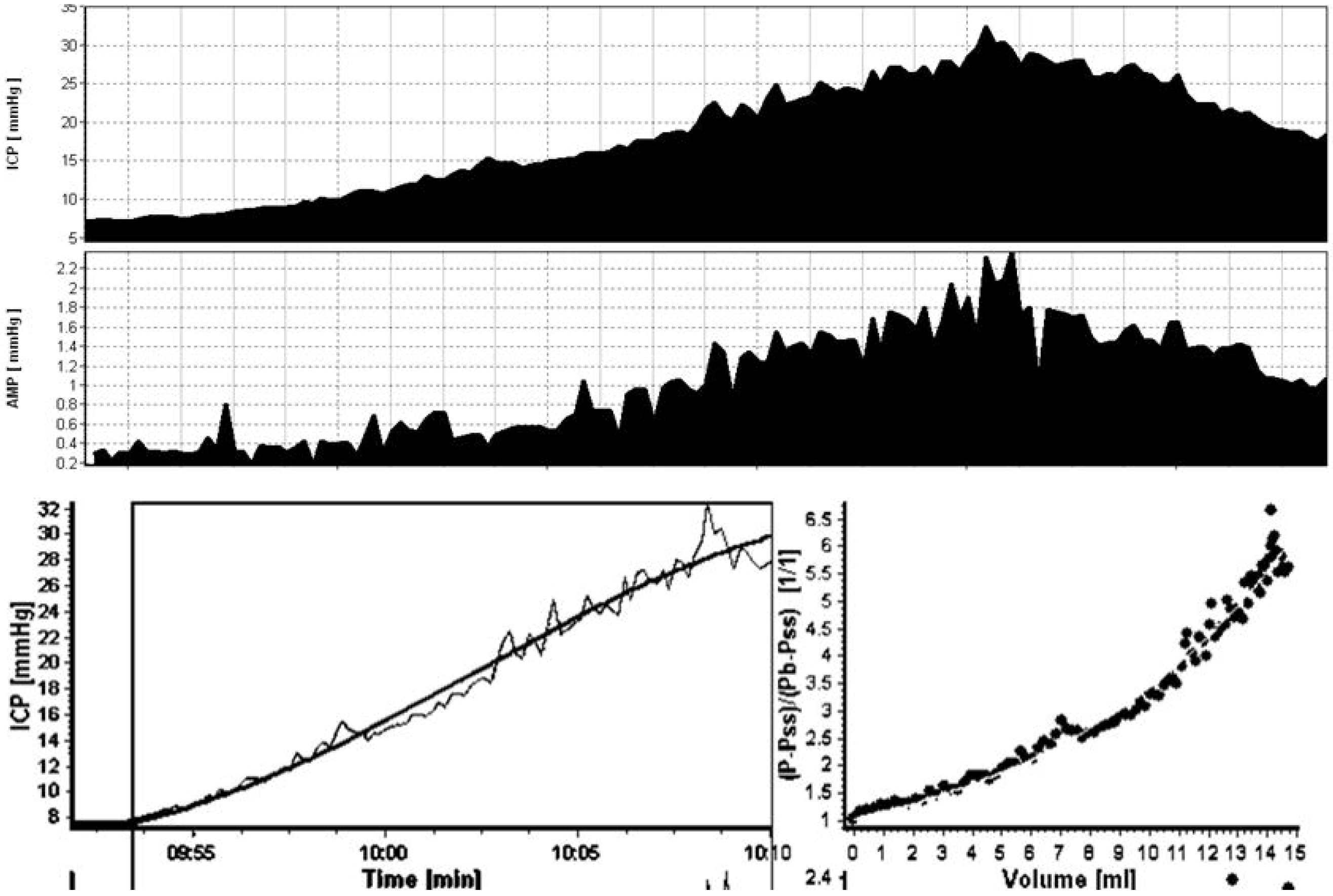
Example of a typical CSP i11fusi.011. study recordi11g mul a11alysis of the test results. Upper panel: Example ofa typical i1ifusion test recording of a patient refened with possible iNPH. ICP (upper area) is monitored at baseline for 5-10 minutes and is gradually increased by Hartmann’s/Ringer’s i,ifusion until a stable plateau). AMP (fundamental amplitude ofICP - lower area) typically follows the increase ofICP. Lower panel; Assessment ofCSF dynamics using CSF models integrated in ICM+ to optimi:e calculation ofRout and other parameters calculated during infusion test. Right curve: the red line (solid,curved line), 1epresents the theoretical model representing the response ofICP to infusion at each time-baseline, during i,ifusion and plateau. The second line represents the calculations pe1formed by the inte1preter ofthe CSF test and which should optimallyfit the theoretical mode. Le.ft c11Jve: The light blue line 1epresents a similar model, where the calculations ofthe user shouldfir the Presswe - Volume curve as proposed by Marmarou. (1974)^19^ and integrated in ICM+. ICP: Intracranial P1ess11Je, AMP: fundamental amplitude of ICP

#### Patient follow-up & outcome assessment

After undergoing a CSF infusion test, the patients are followed up by the clinical team to receive a final clinical diagnosis and also to decide on whether or not to proceed to shunting or ETV. The clinical criteria used up to 2013 to make the final diagnosis have been previously reported [20] and mainly include the Rout (threshold of 13 mmHg/min/ml) but also imaging evidence of the proportion of deep white matter lesions and response to the tap test. Outcome was assessed using a simple scale[23] and based on a combination of improvement primarily and objectively in gait combined with feedback from the patient. Outcome 1 & 2 correspond to clinical improvement for 6 and 3 months after shunting respectively, whereas outcome 3 corresponded to no clinical improvement.

### Statistical Analysis

Data was recorded and processed using ICM+ software. There is ample evidence in the literature for using Rout estimators derived from the computerised CSF infusion test, with values appearing to correlate very well [2, 3, 26, 27]. In a data analysis approach, there are two modes of calculating Rout; static mode, derived from the classic formula (ICP plateau – ICP baseline) (mmHg) / infusion rate (ml/min), and dynamic mode, which derives from optimising this calculation by fitting it to a mathematical model. ICM+ allows for calculation of Rout using both methods. The dynamic calculation is almost always preferred, unless there is no optimal fit between our calculations and the mathematical model, in which case the dynamic calculation of Rout becomes unreliable and as such it is discarded and the static calculation of Rout is used instead.

All statistical analysis was carried out in R software (version 3.3.3). Relationships between variables were sought by calculating Pearson or Spearman product correlation coefficients and p values. Between-group differences (eg different outcome groups) were tested using the Wilcoxon signed rank test or the student t-test, after checking for normality and confirming parametric assumptions. Multiple linear regression was performed to analyse the influence of other variables on the relationships under investigation.

## Results

The study included 229 patients: 137 males and 92 females, male-to-female ratio of approximately 1.5:1. Their age ranged from 36 to 96 years [median age 75years, mean age of the cohort was 70.4 (± 13.82)] at time of the infusion test.

The values calculated for Rout, ICP, AMP and age in male and female subgroups are presented in *table 1* as mean values with standard errors of the mean and level of significance (p-value) when the respective groups are compared.

We found a significant, albeit weak, positive correlation between Rout and ICP (R = 0.17, p=0.0049) as shown in *figure 3A*. On the other hand, there was no correlation between Rout and ICP in the female subgroup, but the correlation was strong and present in the male subgroup (R=0.26, p=0.002). The correlation between Rout and ICP was stronger when investigated by multiple linear regression, which took the influence of age into account (R=0.31, p= 5.935e−06).

**Figure 3.**
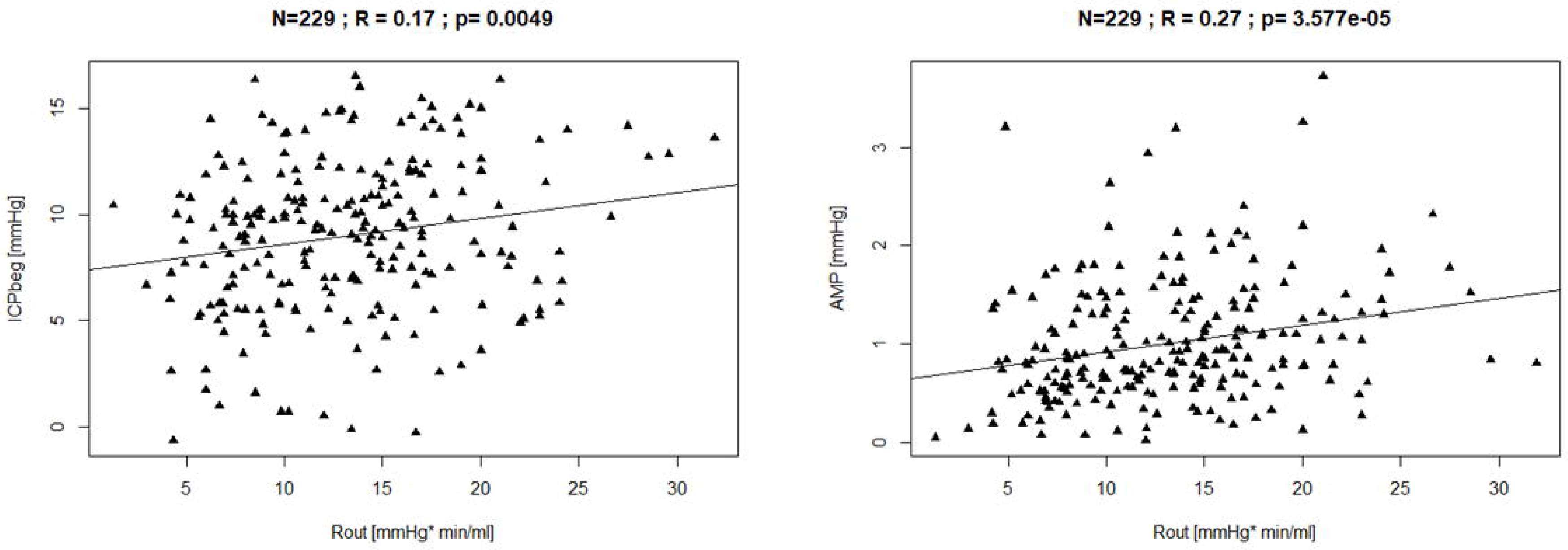
A: Left: Scatter plot showing resistance to CSF outjl.ow versus baseline intracranial pressure. Right: Scatter plot showing resistance to CSF outjl.ow (Rout) versus AMP at baseline. Rout,. resistance to CSF outjl.ow JCPbeg: Baseline intracranial pressure. B: Rout andAMP baseline;

Rout was also positively correlated with AMP (R=0.27, p=3.577e−05), as shown in *figure 3B* and strongly correlated with patients’ age (R=0.16, p=0.01306), as shown in *figure 3C*. The correlation between Rout and age was also absent in the male subgroup, but present and strong in the female subgroup (R=0.33, p=0.001).

**Figure 3B:**
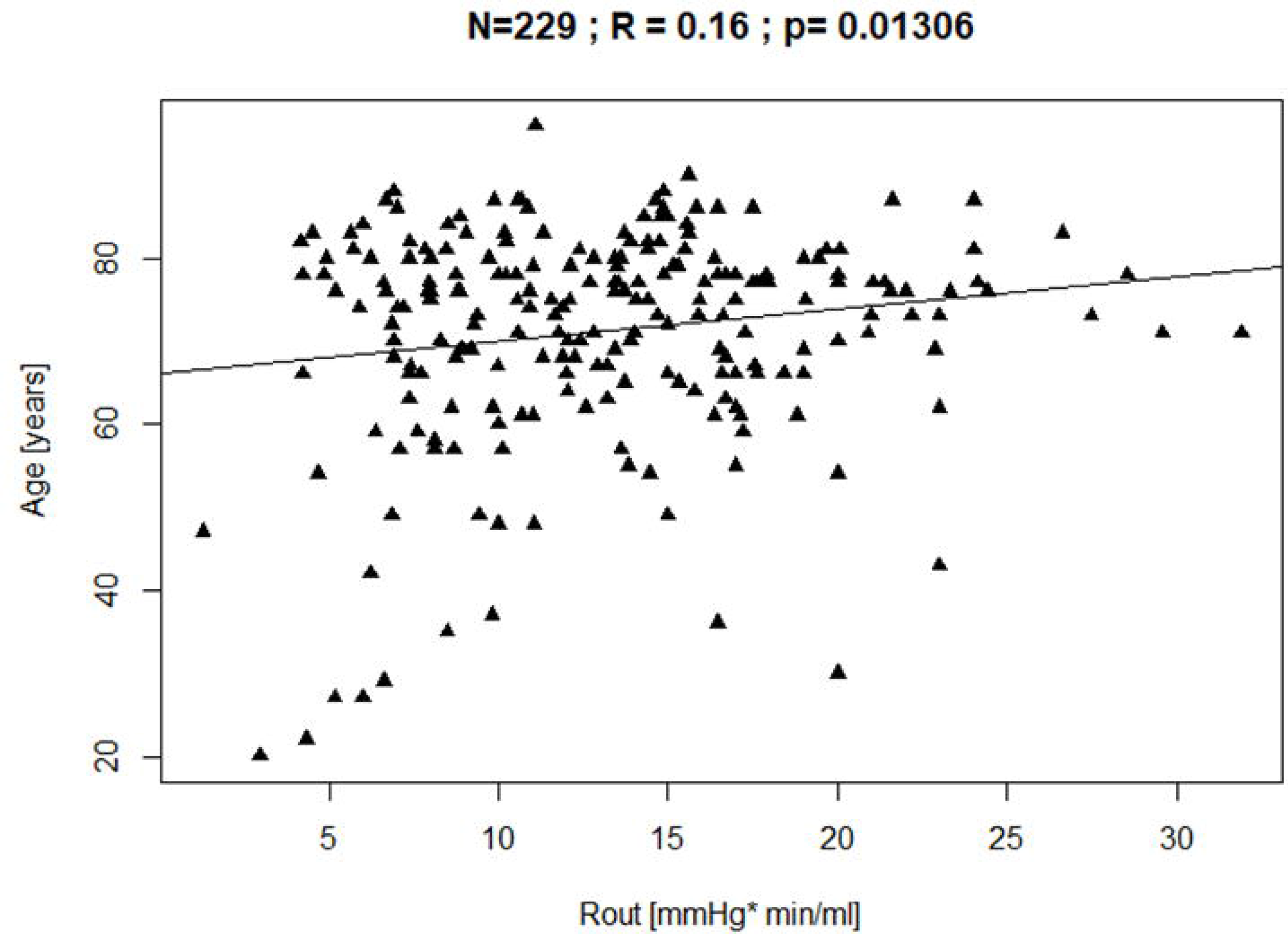
Scatterplot showing resistance to CSF outflow versus Age. Rout,. resistance to CSF outflow.

### Relationship in different diagnosis & outcome groups

Following infusion studies and clinical evaluation,, 149 patients received a final diagnosis of NPH and for 51 did not, whereas the rest of the patients (total of 29) were lost in follow up. 143 were finally shunted or underwent ETV and were available for follow up in our hydrocephalus clinic. 5 patients had ETV and the remaining 138 had shunt insertion. The type of interventional choice was made by the consultant clinician. 119 of those responded well to the CSF diversion (outcome =1 or 2) versus 15 who did not demonstrate any improvement (outcome=3). 9 patients were lost in follow-up.

The correlation between Rout and ICP baseline (ICPbeg) appeared insignificant in the group of patients with a final clinical diagnosis of NPH without considering the effect of age (R=0.1; p=0.0533), but when age was accounted for the correlation became stronger and significant (R=0.3993; p=3.095-06). Therefore, there is a significant correlation between Rout and ICPbeg in the 143 patients with a final diagnosis of NPH provided that we take age into consideration. Furthermore, the relationship was absent in the patients who did not receive an NPH diagnosis. Similarly, the relationship was only present when the interaction between Rout and Age was considered in patients with favourable outcome, that is outcome=1 or outcome=1 or 2 (R=0.43; p = 6.302e−06) while being absent in the non-responders (outcome =3).

## Discussion

The key finding of this study is a weak but significant correlation between Rout and ICP in a large cohort of iNPH patients. This result supports the use of Davson’s equation to model CSF hydrodynamics, however its use without precise knowledge of Pss and CSF formation rate is limited. The positive correlation of Rout and ICP provides evidence supporting Rout as a key disturbance in NPH. It suggests that Rout is implicated in the pathophysiology in NPH and that an elevation of baseline ICP is possible in these cases.

Despite having found a significant correlation, it might not be as strong as would be expected. Possible explanations for this could include variations of pressure in the sagittal sinus, CSF formation rate and those cases where baseline ICP was lower than Pss, where Davson equation is not valid at all. Pressure in the sagittal sinus is considered a constant parameter, determined by central venous pressure[11, 12, 21, 23, 24]. However, despite P_SS_ being constant in one individual, it may vary amongst different individuals. Ekstedt et al found P_SS_ to range from 0.7 to 1.35kPa (5.25 – 10.13 mmHg) in “physiological” individuals[15, 24]. However, it is currently rare to measure P_SS_ directly. This heterogeneity of P_SS_ values in the population could contribute to creating an apparently weaker correlation. Rout was also found to increase with age. Similarly, I_f_ also remains unmeasured in “physiological” individuals and unknown in hydrocephalic individuals[14, 15, 24]. I_f_ is generally reported as stable at 0.35 ml/min, however we are not aware of any studies with reliable I_f_ measurements for hydrocephalic individuals or hydrocephalus animal models.

Rout was found to increase with age, in accordance with previous publications on this existing relationship [20, 23]. Therefore, we considered whether age could act as a confounding variable between Rout and ICP. We carried out a multiple linear regression analysis, which showed a stronger and more statistically significant association between Rout and ICP. This influence shows that the correlation between ICP and Rout has multiple influencing factors. Age is known as one of these factors. For future consideration, it would be interesting to explore formation rate and P_SS_ further, in order to determine the ICP-Rout relationship more holistically. To the best of our knowledge, these parameters and their behaviour in NPH patients has not yet been investigated.

Despite Rout, ICP and AMP not differing between previously defined subgroups (males and females) there were significant differences in the correlation coefficients calculated for these groups. The male and female subgroups each contain a larger number of patients, however sample size is still too small to determine whether the lack of correlation calculated is accurate and further study will be required for confirmation. Furthermore, this is a retrospective study involving patients with a working diagnosis of NPH. A future study with follow up of these patients in respect to improvement after shunt surgery and confirmation of NPH diagnosis could shed more light on our findings.

Alarmingly, the correlation between Rout and ICPbeg demonstrated an even stronger age dependency in the group of patients with a final clinical diagnosis of NPH. This is an interesting finding that could be used to argue in favour of the importance of age-adjustment for Rout calculations in hydrocephalic individuals, to assist NPH diagnosis and shunt prognostication[17–19, 23]. Finally, the similar behaviour of the correlation in the group of patients who positively responded to shunting, that is the strongly age-dependent detection of the correlation, absent in the non-responders, could also have additional value to the new chapter of Rout optimisation and CSF dynamics studies in hydrocephalic patients. The number of patients in the non-responder group is quite low (N=15), however outcome is not the current focus of this study.

## Limitations

We have referred to patients as possible and probable NPH candidates as it is frequently impossible to ascertain whether a patient has NPH, pure NPH or NPH plus other co-morbidities. Unfortunately, there is no gold standard for the manner in which to accurately diagnose NPH, predict shunt response, to decide to shunt or even how to measure response to shunting. Therefore, as there is not always a way to definitively diagnose NPH, it is to be expected that we cannot determine with certainty which of our patients had NPH and which did not. It is possible that some of these patients could have a different diagnosis, resulting in a wide range of Rout values. The influence of different characteristics, such as co-morbidities and duration of symptoms should be taken into account but is not always determined in our patients. Finally, there are very few patients with no response to shunting, so the results from this analysis are subject to the relevant limitations. In the end, the objective truth on the diagnosis of iNPH is unfortunately still open for debate.

As previously reported, the patients referred for infusion studies are referred from a multitude of neurosurgical consultants [20]. Therefore, we are unable to report clinical information, pre-and post-operative assessment of the magnitude of the symptoms and the patients’ improvement in great detail. This is due to a great loss of data that is part of the retrospective nature of the study. However, every patient undergoes thorough investigations by specialists before diagnosis and outcome classifications are made. Furthermore, our simple, 3-level scale does not report in detail the magnitude of improvement of the patients’ symptoms, despite that the patients have been investigated, monitored and followed-up closely in order to determine their management and outcome reflected in our scale.

## Conflict of Interest

All authors certify that they have no affiliations with or involvement in any organization or entity with any financial interest (such as honoraria; educational grants; participation in speakers’ bureaus; membership, employment, consultancies, stock ownership, or other equity interest; and expert testimony or patent-licensing arrangements), or non-financial interest (such as personal or professional relationships, affiliations, knowledge or beliefs) in the subject matter or materials discussed in this manuscript.

### Ethical approval

For this type of study formal consent is not required.

### Informed consent

Informed consent was obtained from all individual participants included in the study.

## Acknowledgements

No acknowledgements

